# Literature mining supports a next-generation modeling approach to predict cellular byproduct secretion

**DOI:** 10.1101/066944

**Authors:** Zachary A. King, Edward J. O’Brien, Adam M. Feist, Bernhard O. Palsson

## Abstract

The metabolic byproducts secreted by growing cells can be easily measured and provide a window into the state of a cell; they have been essential to the development of microbiology^1^, cancer biology^2^, and biotechnology^3^. Progress in computational modeling of cells has made it possible to predict metabolic byproduct secretion with bottom-up reconstructions of metabolic networks. However, owing to a lack of data, it has not been possible to validate these predictions across a wide range of strains and conditions. Through literature mining, we were able to generate a database of *Escherichia coli* strains and their experimentally measured byproduct secretions. We simulated these strains in six historical genome-scale models of *E. coli*, and we report that the predictive power of the models has increased as they have expanded in size and scope. Next-generation models of metabolism and gene expression are even more capable than previous models, but parameterization poses new challenges.

## 1 Introduction

All cells secrete metabolic byproducts in the course of growing and producing energy, and these byproducts play important roles in the study of biological systems. Byproducts are a readout of the cellular state; lactate excretion, for instance, is characteristic of tumor cell growth^2,4^. Byproducts can be engineered for bioproduction of commodity chemicals and biofuels^5–7^. And byproducts of yeast fermentation – including ethanol – are responsible for the most popular beverages in human history^8^. With the critical roles played by metabolic byproducts in disease and biotechnology, it is of great interest to be able to predict the byproducts that a cell will secrete under a specific condition. However, no published study has assessed whether existing computational methods are able to predict metabolic byproducts for a range of strains and conditions.

Computational models have been shown to correctly predict byproduct secretion under common laboratory conditions. During aerobic growth, the model bacterium *Escherichia coli* oxidizes substrate molecules to secrete CO2 and water; during anaerobic fermentation, *E. coli* secretes mixed-acid fermentation products (ethanol, acetate, formate, D-lactate, and succinate)^9^. Genome-scale models (GEMs) and constraint-based reconstruction and analysis (COBRA) methods rely on knowledge of the metabolic network and mass-balance during steady state growth to predict the optimal distribution of metabolic flux for growth^10^. GEMs have been shown to be able to predict *E. coli* byproduct secretions in certain cases^11,12^. In the context of GEMs, the byproducts that must be secreted for optimal growth are called *growth-coupled*, and computational methods have been developed to predict and engineer growth-coupled chemical production^13–15^. However, few experimental studies have followed from the computational method development (among them:^12,16^), so it is unclear how these methods would scale up to a wide variety of strains and conditions.

Next-generation GEMs of metabolism and gene expression (called ME-models^17–19^) are now available; ME-models predict the composition of the entire proteome of a cell. In contrast, GEMs of metabolism (M-models) predict only the reaction fluxes in a metabolic network^18^. One new capability of ME-models is the ability to predict the bacterial Warburg effect, the tendency of bacteria to secrete acetate during aerobic growth in the presence of excess substrate^4,20^. In ME-models, the limitations of ribosome efficiency lead to low-yield metabolic approaches like acetate secretion^17^. The same effect can be seen in smaller-scale growth models and is supported by phenotypic data^4,20^. Whether ME-models can correctly predict byproduct secretion for other conditions is not currently known.

High-quality genotypic and phenotypic data are required to test any model predictions, and such data have not been available for the study of byproduct secretion. The present study takes a novel approach by mining the research literature for examples of engineered strains of *E. coli* with diverse byproduct secretion mixtures. We collected 73 papers reporting a total of 89 strains of *E. coli* that have a wide range of gene knockouts, heterologous pathways, and growth conditions, and we simulated these paired genotype-phenotype data in 6 historical GEMs of *E. coli*, including the next-generation ME-model. We find that GEMs have been improving in their ability to recapitulate measured byproducts from experimental studies as the models have increased in size and scope. We explore the possible reasons for incorrect predictions and provide insights into the challenges of simulating byproduct secretion for any growing cell.

## 2 Results

### Literature mining provides a diverse set of strains and phenotypes

An impressive body of data on *E. coli* byproduct secretion can be found in the peer reviewed literature (Fig. 1). We generated a bibliomic database using a workflow for identifying relevant papers, extracting data, and performing quality assessment (Fig. S1). Each paper in the database reported a strain design of *E. coli* in which the fermentation pathways were engineered to force the cell to secrete a target molecule (Fig. 2). The bibliomic database includes the gene knockouts, heterologous pathway descriptions, substrate conditions, oxygen availability, and the parent cell line for each strain (Supplementary Data 1). It is difficult to extract and normalize quantitative measures of byproduct secretion from the literature. Instead, we recorded the molecule that was targeted for overproduction in the study, and we confirmed that this byproduct was the major secretion product in each case (see Methods). The bibliomic database contains 73 papers and 89 strains of *E. coli*; this is approximately 20% of all papers on metabolic engineering of *E. coli* collected in the LASER database^21^.

**Figure 1:**
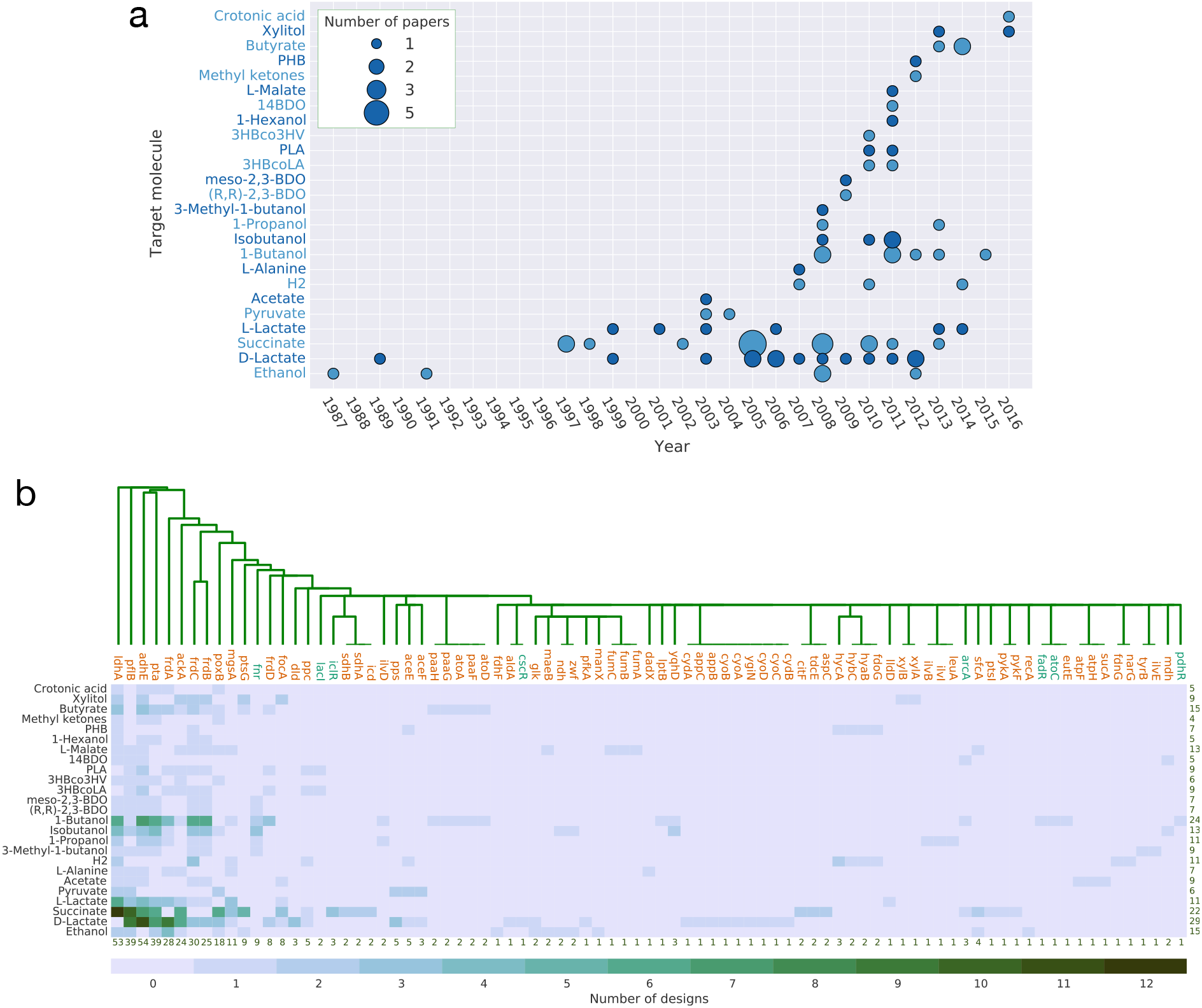
The bibliomic database. (a) The number of papers published for each target molecule over time. (b) The individual genes that have been knocked out for each target molecule. The sums across the bottom indicate the total number of designs that include a given gene, and the sums across the right indicate the total number of unique genes knocked out for a given target molecule. These common knockouts remove the routes to the native fermentation products acetate (*pta, ackA, pflB*), ethanol (*adhE, pflB*), formate (pflB), D-lactate (*ldhA*), and succinate (*frdABCD*). These knockouts represent a common strategy where the highest-yield fermentation pathways are knocked out, one by one, until the target pathway becomes the optimal route for balancing the redox state of the cell.

**Figure 2:**
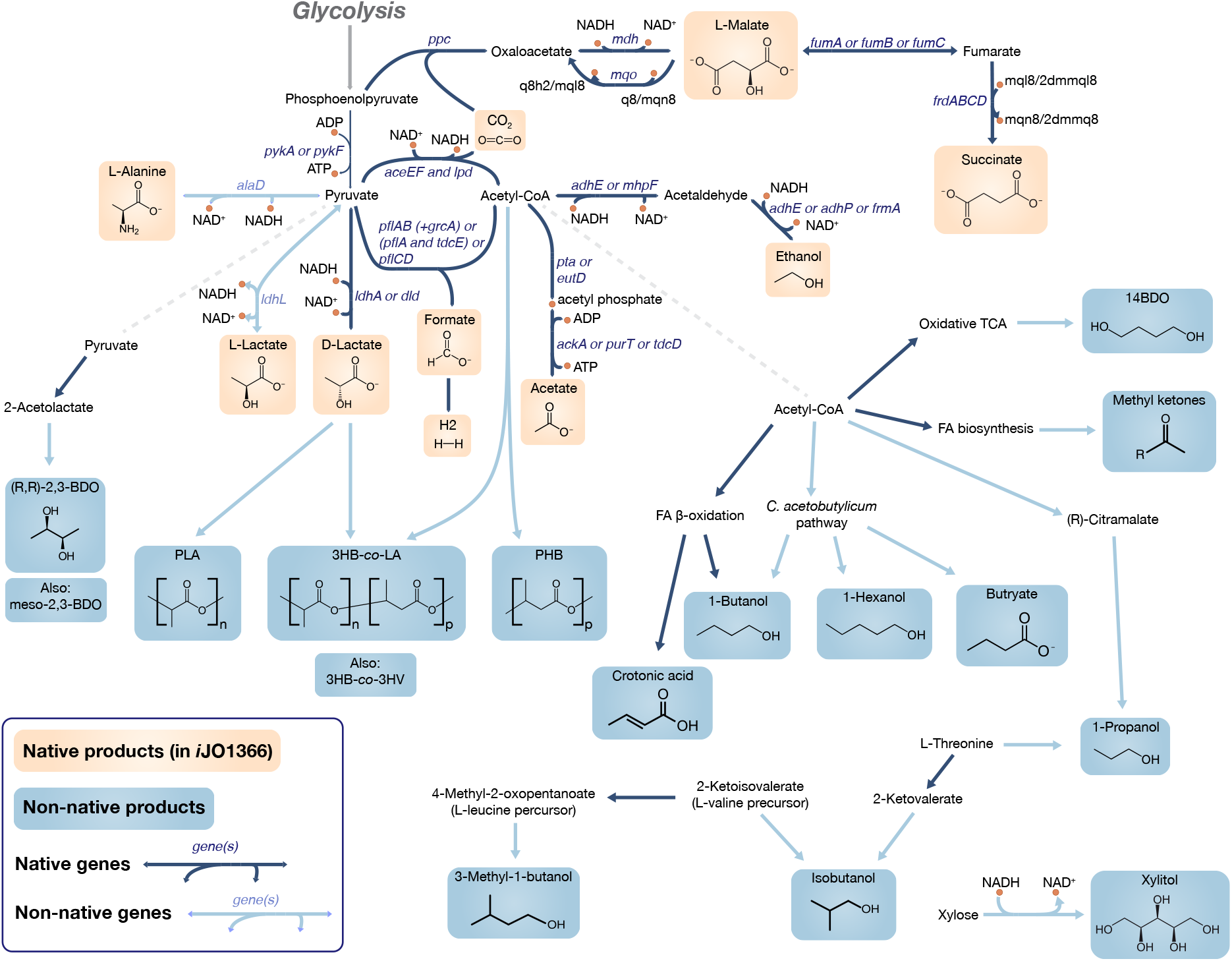
The engineered fermentation pathways in *E. coli*. All the engineering pathways in the bibliomic database are shown, along with their metabolic precursors. Native products (yellow) are those that appear in the genome-scale model iJO1366. Native pathways in iJO1366 (dark blue arrows) and non-native pathways (light blue arrows) are also differentiated.

The strains in the bibliomic database were simulated in six GEMs of *E. coli* (Table 1). The models have increased in size and complexity over the past decade; they include five M-models and one ME-model that includes 1,683 genes and accounts for 80% of the proteome by mass^17,18^. Gene knockouts, heterologous pathways, and environmental conditions from the bibliomic database were recreated in each of the GEMs. For each strain, flux balance analysis (FBA)^22^ was used to find the predicted growth rate and the growth-coupled yield, the carbon yield of a compound at the maximum growth rate. The analysis began with two comparisons between the bibliomic database and the simulations: (1) whether the strain grew in a given environment and (2) whether the simulation predicted growth-coupled secretion of the target byproduct from the study.

**Table 1:** The increasing size and scope of genome-scale models of *E. coli*.

| Model | Genes | Reactions | Metabolites/Components | Year (Reference) |
| --- | --- | --- | --- | --- |
| Core model | 137 | 95 | 72 | 2006<sup>50</sup> |
| *iJR904* | 904 | 1075 | 761 | 2003<sup>51</sup> |
| *iAF1260* | 1,260 | 2,382 | 1,668 | 2007<sup>52</sup> |
| *iAF1260b* | 1,260 | 2,388 | 1,668 | 2010<sup>14</sup> |
| *iJO1366* | 1,366 | 2,583 | 1,805 | 2011<sup>45</sup> |
| *iOL1650-ME* | 1,683 | 12,009 | 6,563 | 2013<sup>17</sup> |

The predictive power of GEMs has generally increased over time, with the increasing size and scope of the models. New GEMs provide better predictions of growth-coupled secretion compared to their predecessors (“Model accuracy” in Fig. 3). In order to understand the reasons for this trend, we designed a computational approach to categorize cases of incorrect prediction. Exhaustive search and parameter sampling were employed in the M- and ME-models, respectively, to determine what changes to the modeling approach might lead to *in silico* secretion of the target byproduct (see Methods). These categories provide insights into the general challenges of modeling byproduct secretion.

**Figure 3:**
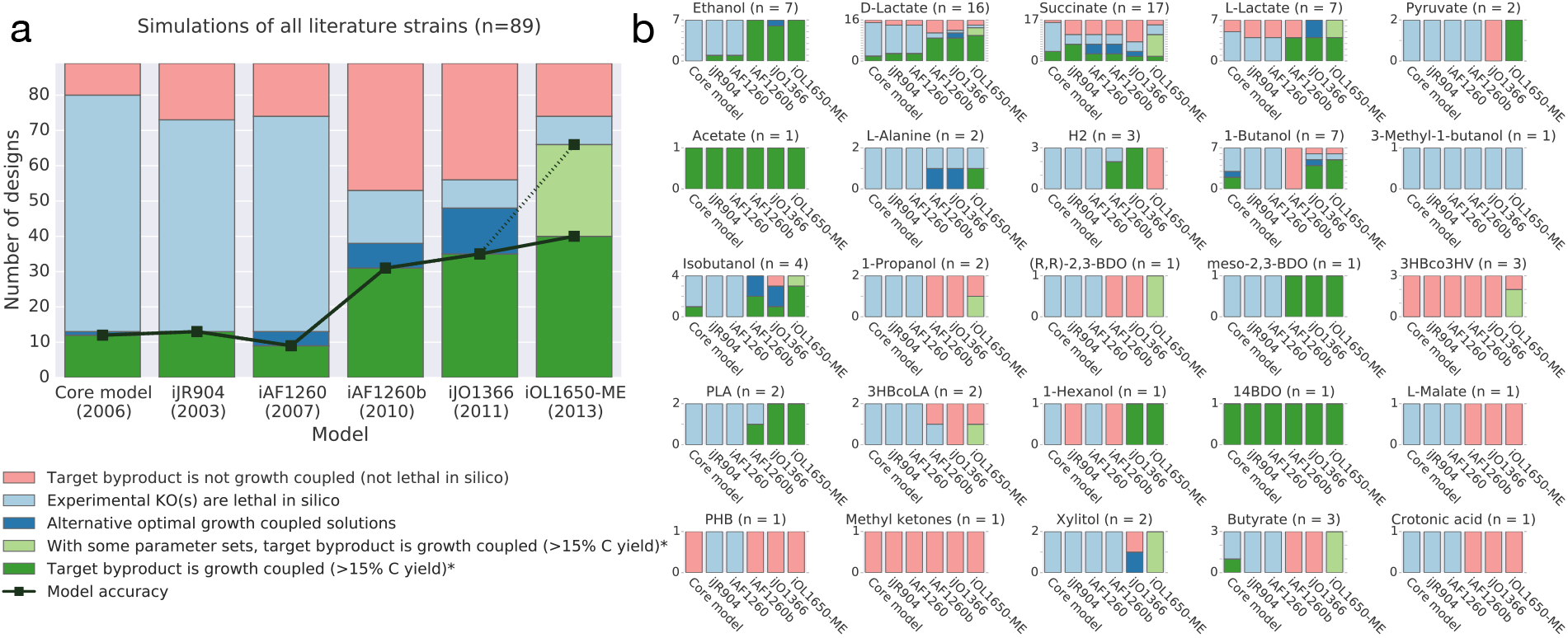
Simulations of the bibliomic dataset in E. coli GEMs. (a) The 89 strains in the bibliomic database were simulated in six GEMs of *E. coli*, and the incorrect predictions were categorized to suggest a reasons for the errors. The solid line signifies that the experimentally observed target byproduct is growth-coupled in the model. The dashed line represents the possibility of improving predictions in the ME-model by correctly determining the kinetic parameters (*k_eff_*S). (b) The categories separated according to the target molecule.

### Genome-scale models do not differentiate between isozymes

Isozymes are common in metabolic networks, and they are represented in M-models, but their diverse regulatory and catalytic properties lead to a broad and complex set of challenges for metabolic modeling. Reactions are often catalyzed by a major isozyme that is responsible for most catalysis, while minor isozymes are also present in the cell but have a smaller role (they may not be expressed or have less-favorable kinetics)^23^; recent progress in studying enzyme promiscuity and underground metabolism suggests that isozymes are even more widespread than previously thought^24^. Many experimental studies report gene knockouts of major isozymes that decrease the activity of the associated reaction significantly, enough so that the minor isozymes can be ignored (e.g. removing *ldhA* and ignoring *dld*^25–27^). However, M-models do not distinguish between major and minor isozymes, so these cases are incorrectly predicted in the model; the minor isozyme catalyzes the reaction *in silico*, and the *in silico* gene knockout of the major isozyme has no effect. Therefore, to simulate byproduct secretion for real-world experiments, it was necessary to employ a “greedy knockout” strategy in which all reactions associated with a gene knockout are disabled, even if minor isozymes might be present (Fig. 4a).

**Figure 4:**
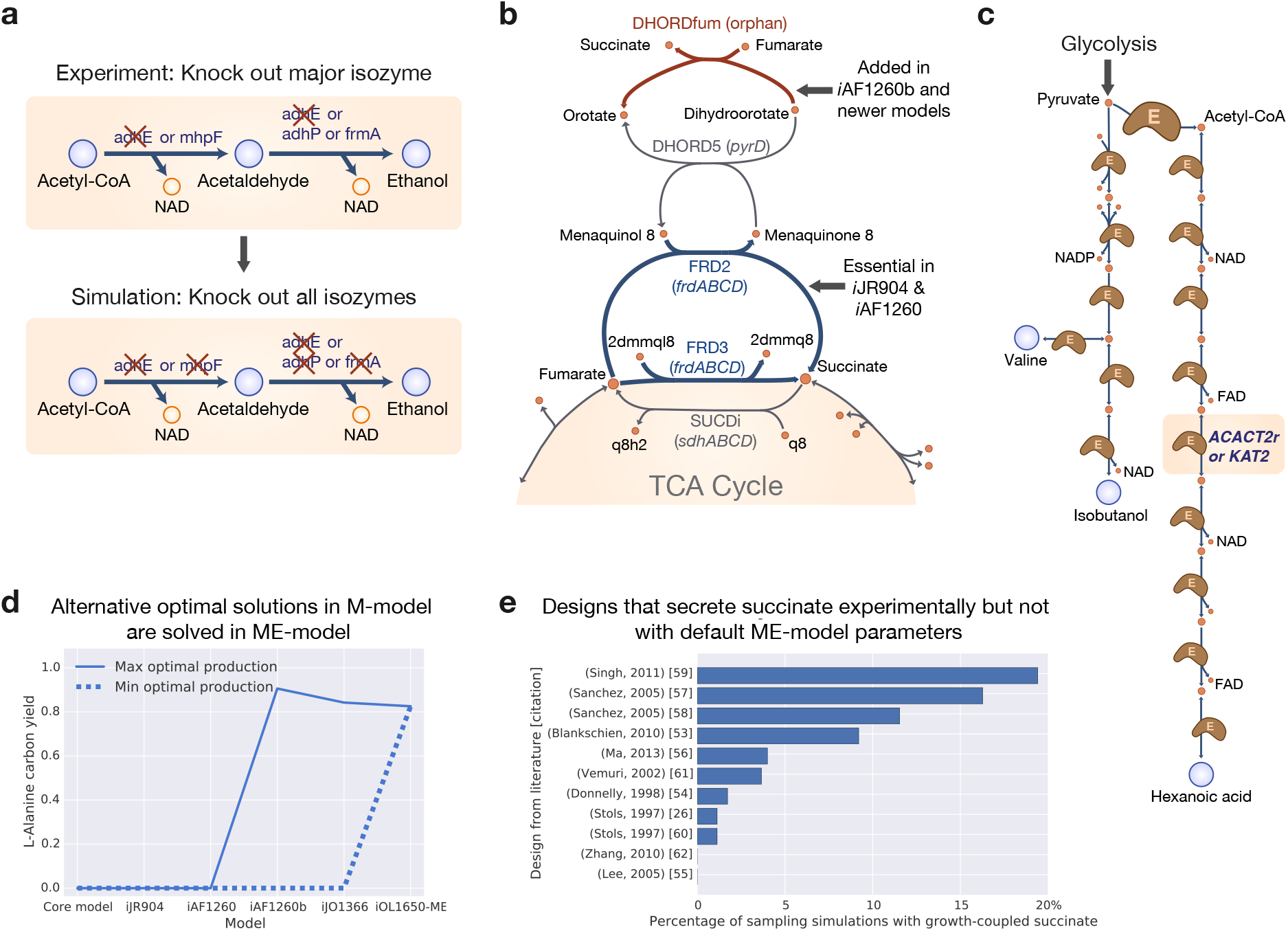
Comparing simulations with experiments. All modeling approaches have failure modes, and comparing model predictions to experimental results allows these failure modes to be analyzed. (a) A “greedy knockout” strategy is necessary to contend with major and minor isozymes that are difficult to simulate in GEMs. (b) The genes in the *frd* operon are responsible for most of the incorrect predictions of cell death in *i*JR904 and *i*AF1260. This error was fixed in *i*AF1260b and later models with the addition of the reaction DHORDfum. (c) For an isobutanol design, the ME model correctly predicts isobutanol secretion in preference to hexanoic acid secretion because the hexanoic acid pathway has greater protein cost^34,35^. (d) Alternative optimal phenotypes appear in M-models when two pathways have equivalent stoichiometries, as in this example for L-alanine secretion. ME-models explicitly account for the cost of producing pathway enzymes, so the shorter L-alanine production pathway is optimal in ME-models. (e) Succinate secretion is difficult to predict using existing GEMs, but an ensemble of ME-models with sampled kinetic parameters demonstrates that for certain parameter sets succinate secretion is correctly predicted.

There are exceptions where greedy knockouts are not appropriate. For example, the alanine racemase activity of isozymes *alrR* and *dadX* is necessary for *in silico* growth, so applying the greedy knockout strategy to the reported strain that has a knockout of *alrR* leads to a prediction of cell death^28^. In other words, this strain can not be correctly simulated by M-models with or without the greedy knockout strategy. This issue can only be addressed through continued development of genome-scale modeling methods to address regulation, kinetics, allosteric inhibition, and the many biophysical properties that differentiate isozymes. Furthermore, ME-models can potentially select the appropriate enzyme based on protein cost, but ME-models do not include regulatory effects that often are responsible for the distinction between major and minor isozymes, so greedy knockouts are still generally required. In this study, the greedy knockout approach was sufficient to correctly simulate most of the gene knockouts in the bibliomic database.

### Larger models solve false predictions of cell death

Every strain in the bibliomic database was able to grow in the published experimental studies, but many simulations of these strains in early GEMs resulted in predictions of no growth (defined as *in silico* specific growth rate less than 0.005 hr^−1^). These incorrect predictions have decreased as the GEMs have increased in size and scope (“Experimental KO(s) are lethal *in silico*” in Fig. 3). In most cases, the reason for the improved prediction is that the more comprehensive GEMs include a pathway that can rescue an essential cellular function when another important pathway is disabled by gene knockouts. In the five *E. coli* M-models, the lethal genotypes were analyzed by exhaustively searching for the minimal combinations of reactions that lead to *in silico* cell death (Fig. S2).

The biggest improvement in modeling the strains in the bibliomic database can be attributed to a single reaction. The models *i*JR904 and *i*AF1260 incorrectly predict that fumarate reductase (FRD, frd) is essential under anaerobic conditions, and 63% of the designs in the bibliomic database include a knockout in the *frd* operon (see the large jump from *i*AF1260 to *i*AF1260b in Fig. 3a). These incorrect predictions were corrected in *i*AF1260b and later GEMs with the inclusion of a new reaction (DHORDfum) that rescues growth when FRD is removed (Fig. 4b). However, there is no experimental evidence to support the presence of the DHORDfum reaction. So why does this reaction exist in the models, and why does it improve predictions?

One explanation is that the DHORDfum reaction does not take place in the cell, and, instead, succinate dehydrogenase (SUCDi, *sdh*) acts in the reverse direction to rescue conversion of fumarate to succinate; this has actually been shown experimentally^29^. Thus, the evidence supports removing DHORDfum from the models and making SUCDi reversible. However, this change introduces the challenges associated with modeling isozymes for the activity catalyzed by *frd* and *sdh*, so the presence of DHORDfum has served as a convenient hack for modeling *E. coli*.

### Simulations suggest that some strains have room to evolve

When the experimental observations of byproduct secretion disagree with predictions, another possible explanation is that the experimental strain could evolve to grow faster by adopting the byproduct secretion strategy predicted by the model (“Target byproduct is not growth-coupled” in Fig. 33). FBA simulations predict the metabolic state of a cell that is operating close to optimal growth; GEMs are powerful for predicting cellular behavior precisely because fast growing cells often adopt a near-optimal strategy for growth^30,31^. Thus, some of the disagreement between observation and prediction might be caused, not by model errors, but rather by an assumption of the modeling approach (the optimality assumption). This hypothesis can be tested through laboratory evolution by passing the strain repeatedly^12^. (The process is also called serial passage, metabolic evolution, growth rescue, or adaptive laboratory evolution (ALE).) Laboratory evolution was used in 14 studies (19 strains) in the bibliomic database to improve byproduct secretion, and the predictive power of the model is greater for these cases than for the bibliomic database in general (Fig. S3). This supports the hypothesis that FBA predicts byproduct secretions that are not correct for the reported strains but would be correct if the strains were evolved through growth selection.

### Next-generation ME-models improve predictions but require parameterization

ME-models expand upon M-models by explicitly accounting for all of the biochemical reactions in the gene expression machinery of the cell (including transcription and translation)^17,18^. To include protein production in the ME-model, one must estimate the turnover rate of each enzyme (*k_eff_*) that determines how many active proteins must be present to convert one set of reactants to products in a given time. ME-model simulations used a set of experimentally validated kinetic parameters from a recent study^32^. For high-flux reactions, the *k_eff_*S were shown to be consistent across four growth conditions. However, it is still possible for *k_eff_*S to change between conditions, depending on metabolite concentrations and other variables (they range between 0 and *k_cat_*). Therefore, we sampled *k_eff_*S in the ME-model to generate an ensemble of models for each strain that was not growth-coupled with default parameters (see Methods). We found that 26 / 41 strains in this set could be growth-coupled in the ME-model with at least one model in the ensemble, including 9/11 designs for succinate production (Fig. 4e). Addressing kinetic parameters will have to be a part of ME-model development going forward, and this should lead to better predictions of byproduct secretion.

The protein costs associated with metabolic pathways in the ME-model also solve another failure mode in M-models: alternative optimal solutions. Alternative optimal solutions occur in M-models when two metabolic states lead to the same growth rate, and this common failure mode has been solved with next-generation ME-models (“Alternative optimal growth-coupled solutions” in Fig. 3)^33^. In ME-model simulations, each pathway has specific enzyme costs that must be precisely allocated using cellular resources. Therefore, pathways with the same metabolic contribution to cellular growth (e.g. same ATP production and redox balance) that are equivalent in the M-model have different proteomic costs in the ME-model. In all cases, this failure mode of M-models disappear in ME-model predictions (with one example provided in Fig. 4d).

In addition to removing alternative optimal solutions, the proteomic pathway costs in the ME-model can address challenges of encoding reversibility in the M-model. As an example, the production of isobutanol using a 2-keto acid based pathway was recently demonstrated^34,35^, and the optimal *in silico* phenotype of this production strain varies between models of *E. coli* (Fig. 3b). *i*AF1260b correctly predicts the production of isobutanol as the optimal fermentation product; in contrast, *i*JO1366 predicts that hexanoic acid, a 6-carbon intermediate in the *β*-oxidation cycle, is the preferred product. This difference can be traced to the thermodynamic reversibility of the thiolase reaction in the second round of the reversed *β*-oxidation cycle – it is irreversible in iAF1260 (KAT2) and reversible in *i*JO1366 (ACACT2r) (Fig. 4c). The reversibility in *i*JO1366 is in line with experimental evidence^36^, but it also leads to the seemingly incorrect prediction of hexanoic acid secretion. The ME-model suggests that the incorrect prediction of hexanoic acid secretion by *i*JO1366 is not so much a matter of thermodynamics as a matter of pathway length and thus proteomic cost. When the cost of producing enzymes for metabolic pathways is incorporated into genome-scale models, long pathways like the hexanoic acid production route through *β*-oxidation carry a greater cost than the shorter 2-keto acid route to isobutanol. This case shows the power of a constraint-based modeling approach: Properly encoding reversibility in M-models has been a long-standing challenge, so the ME-model applies a completely different constraint (pathway cost) that makes the reversibility of *β*-oxidation unimportant for correct predictions.

## 3 Discussion

As cellular models become larger and more complicated, the datasets used to validate them must also grow. This study presents a novel approach to model validation based on literature mining. In spite of the uneven quality of literature data, this approach was capable of generating important insights into the abilities of GEMs to predict byproduct secretion. Higher-quality data would enable an even more thorough model validation, and there is a great need in systems biology for standardizing genotype-phenotype datasets. Standards for storing phenotypic data have been discussed^37,38^, and it is essential that progress be made.

There are a few challenges that will have to be addressed to scale these methods to larger and more complicated systems. First, many data points in the bibliomic database cannot be modeled in existing GEMs. For instance, regulatory knockouts are not in the scope of M- and ME-models, so they were ignored in this study. The correct predictions of strains in the bibliomic database draw largely from the concept of redox balance in the cell (NAD(P)H produced during glycolysis must be consumed by fermentation pathways), and extending prediction of byproduct secretion to other applications where redox balance is not the driving phenomenon may require further development of the modeling methods. However, constraint-based modeling methods are generally extensible, as we have seen with the development and implementation of ME-models. Exploration of constraint based approaches to other subsystems – including protein structures, membrane translocation, and regulation – are under way^39^.

Second, strains modeled using GEMs and FBA must be operating close to an optimal growth state. Understanding the byproduct secretion of strains that are not growing rapidly will require research into other objective functions that could make the models predictive for strains that are not optimizing for growth^40,41^. On the other hand, the optimality assumption of FBA offers an advantage: GEMs and laboratory evolution can be used together for systematic optimization of microorganisms^12,16^.

Finally, the extension of these methods to larger and more complex organisms, such as tumor cells, will require rigorous development and assessment of GEMs. This study provides an example of validating model predictions using genotype-phenotype data mined from the literature. The collection of these data will need to be scaled up to validate larger and more complex models. All cells have the same basic features that include gene expression, metabolism, and, by necessity, byproduct secretion; with targeted validation studies, we can feel increasingly confident in our ability to model and understand them.

## 4 Methods

### Literature mining

A literature mining search was performed to identify all papers reporting the construction of a cell factory strain of *E. coli* for the production of a fermentation product. A workflow was developed (Fig. S1), hundreds of papers were collected, and 73 were included in the bibliomic database based on their matching the following criteria:

- Utilized a strain of *E. coli*.
- Modified the strain for production of a native or heterologous metabolite.
- Removed alternative fermentation pathways using gene knockouts.

Metadata were collected from each paper, including the target production molecule, whether simulations were performed to identify knockouts, the parent *E. coli* strain, the genetic additions and deletions, the aerobicity and carbon sources during fermentation experiments, whether laboratory evolution was performed, and (when possible) the measured fermentation profile of the engineered strain.

A single target molecule was selected for each experiment, even though in some cases a mixture of products was reported. When papers reported mixtures of hydrogen or formate with a coproduct, the coproduct was considered the target molecule.

### Simulations

To simulate reported designs, the gene knockouts were implemented *in silico* using a “greedy knockout” strategy. For each gene that was knocked out experimentally, all reactions associated with that gene in the metabolic model are turned off. The alternative strategy is to evaluate the gene-protein-reaction (GPR) rules for each reaction in turn, to determine whether the reaction is turned off or remains unchanged; however, as discussed in the text, only the “greedy knockout” approach was able to correctly simulate strains in the bibliomic database.

For all non-native genes reported in the papers, pathways were reconstructed by creating *in silico* reactions corresponding to the genes used in these experiments. For transport reactions, transport was assumed to be non-energy-coupled unless otherwise specified in the iJO1366 reconstruction or in the literature.

Polymer production must be considered separately from ordinary metabolite secretions. To simulate these strains, the production of the monomer was optimized. It is unclear whether polymers such as polylactic acid (PLA) would be growth coupled. The PHA synthase is not energy coupled^42^, so an equilibrium between monomer and polymer would probably be achieved in the optimal state (this has been shown for soluble heteroglycans^43^). However, by upregulating the PHA synthase in an strain optimized for monomer production, one can use the growth-coupling effect to perform much of the strain optimization. Thus, growth-coupling of the monomer is of interest.

Five M-models and one ME-model of *E. coli* K-12 MG1655 were used for the simulations in this work. The M-models were collected from the BiGG Models database^44^, and they were used as reported in their respective publications (Table 1). As described previously, the *i*JO1366 oxidative stress reactions CAT, SPODM, and SPODMpp and the FHL reaction were constrained to zero^45^. A new software implementation of the ME model iOL1650-ME was used. Pathway diagrams were generated using Escher^46^, and COBRA simulations were performed with COBRApy^47^.

For M-model simulations, the substrate uptake rates (SURs) for the solitary carbon substrates in each simulation were constrained to a maximum uptake rate of 10 mmol gDW^−1^ hr^−1^. The oxygen uptake rates were constrained to 0 for anaerobic conditions and 20 mmol gDW^−1^ hr^−1^ for aerobic conditions. For ME-model simulations, SURs were left unbounded and the ME-model optimization procedure chose optimal SURs. If LB or yeast extract was present in the medium, the simulations were still performed with an *in silico* minimal media based on the assumption that cells will preferentially consume glucose before more-complex carbon sources; however, if this approximation led to a lethal phenotype in *i*JO1366, then supplementations known to exist in rich media were added to alleviate the lethal phenotype. Microaerobic designs were assumed to be anaerobic because it has been observed that even under aerobic conditions the anaerobic physiology contributes to fermentation^48^.

FBA was used to find the maximum and minimum secretion of each metabolite in the network when the growth rate is near its maximum (within 0.01%)^22^. The key outputs of these simulations are *predicted growth rate* – the flux through the biomass objective function – and the *growth-coupled yield* – the minimum carbon flux through the target molecule exchange reaction at the maximum growth rate

### Parameter sampling

Parameter sampling in the ME-model was employed to determine the sensitivity of ME-model simulations to *k_eff_* values. For each sampling simulation, an ensemble of 200 models was generated with *k_eff_* values selected randomly from a lognormal distribution of possible *k_cat_*s. The distribution was determined from a collection of all *k_cat_*s in the BRENDA enzyme database (*μ* = 2.48 and *σ* = 3.29)^49^.

### Failure model categorization

Growth-coupling was defined as secretion of the target molecule with greater than 15% carbon yield or, for hydrogen production, greater than 2 mmol gDW^−1^ hr^−1^. Lethal phenotypes were defined as having an *in silico* growth rate below 0.005 hr^−1^. Alternative optima were identified by finding designs whose maximum secretions were above the threshold for growth coupling but whose minimum secretions were below this threshold.

## 5 Acknowledgments

We would like to thank Gabriela I. Guzmán and Joshua A. Lerman for their guidance and suggestions during the course of this project. Funding for this work was provided by the National Science Foundation Graduate Research Fellowship [DGE-1144086 to Z.A.K.] and by the Novo Nordisk Foundation through the Center for Biosustainability at the Technical University of Denmark [NNF16CC0021858].

## 6 Author contributions

Z.A.K. ran the analysis. Z.A.K., E.J.O., A.F.M., and B.O.P. designed the study and wrote the manuscript.

## 11 Extended Data

**Figure S1:**
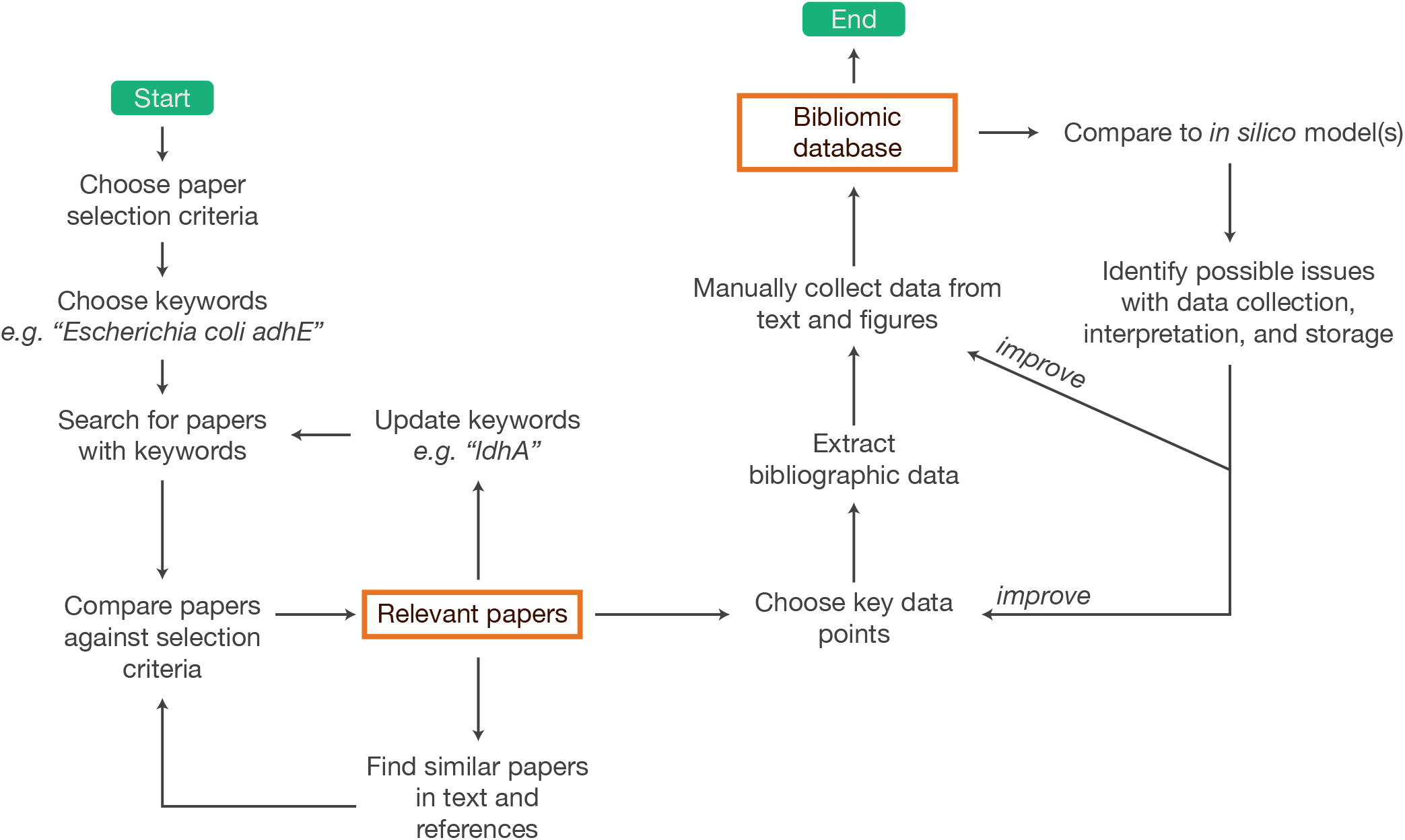
A workflow for generating a bibliomic dataset through literature mining.

**Figure S2:**
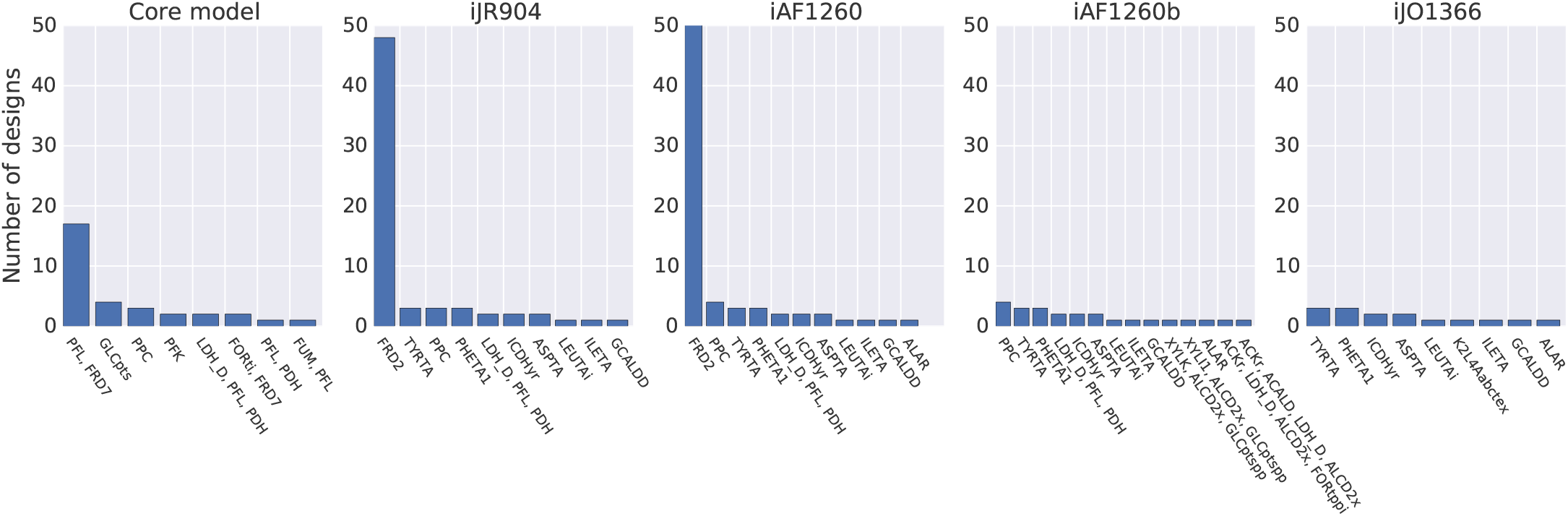
The combinations of reaction knockouts that are lethal in the M-models after simulating the bibliomic database.

**Figure S3:**
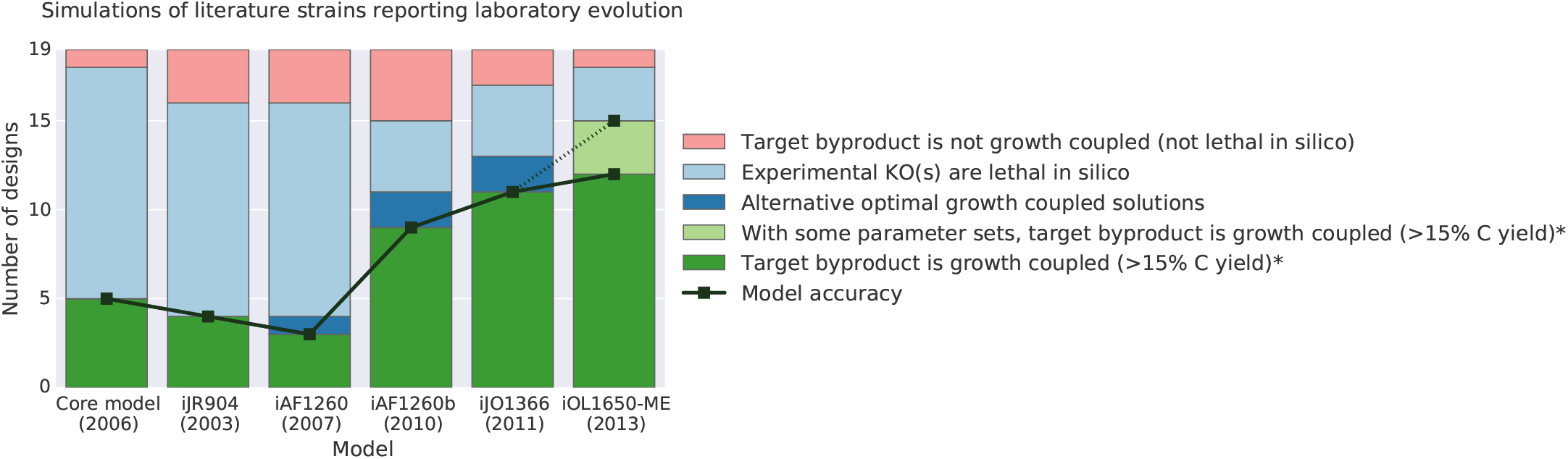
Comparing simulations with experiments for studies reporting laboratory evolution.

## 12 Supplementary Data

Supplementary Data 1: The complete bibliomic database (tab-separated text file).

## References

1. Barnett, J. A. Beginnings of microbiology and biochemistry: the contribution of yeast research. Microbiology 149, 557–567 (2003).

2. Hanahan, D. & Weinberg, R. A. Hallmarks of cancer: the next generation. Cell 144, 646–674 (2011).

3. Demain, A. L. History of industrial biotechnology. In Soetaert, W. & Vandamme, E. J. (eds.) Industrial Biotechnology: Sustainable Growth and Economic Success, chap. 1, 17–77 (Wiley-VCH, Weinheim, Germany, 2010).

4. Basan, M. et al. Overflow metabolism in bacteria results from efficient proteome allocation for energy biogenesis. Nature (2015).

5. Lee, S. Y. & Kim, H. U. Systems strategies for developing industrial microbial strains. Nat. Biotechnol. 33, 1061–1072 (2015).

6. Zhang, F., Rodriguez, S. & Keasling, J. D. Metabolic engineering of microbial pathways for advanced biofuels production. Curr. Opin. Biotechnol. 22, 775–783 (2011).

7. Chubukov, V., Mukhopadhyay, A., Petzold, C. J., Keasling, J. D. & Martín, H. G. Synthetic and systems biology for microbial production of commodity chemicals. npj Systems Biology and Applications 2, 16009 (2016).

8. Piškur, J., Rozpedowska, E., Polakova, S., Merico, A. & Compagno, C. How did saccha-romyces evolve to become a good brewer? Trends Genet. 22, 183–186 (2006).

9. David P. Clark. The fermentation pathways of *Escherichia coli*. FEMS Microbiol. Rev. 63, 223–234 (1989).

10. Bordbar, A., Monk, J. M., King, Z. A. & Palsson, B. O. Constraint-based models predict metabolic and associated cellular functions. Nat. Rev. Genet. 15, 107–120 (2014).

11. Varma, A., Boesch, B. W. & Palsson, B. O. Stoichiometric interpretation of *Escherichia coli* glucose catabolism under various oxygenation rates. Appl. Environ. Microbiol. 59, 2465–2473 (1993).

12. Fong, S. S. et al. In silico design and adaptive evolution of *Escherichia coli* for production of lactic acid. Biotechnol. Bioeng. 91, 643–648 (2005).

13. Burgard, A. P., Pharkya, P. & Maranas, C. D. Optknock: a bilevel programming framework for identifying gene knockout strategies for microbial strain optimization. Biotechnol. Bioeng. 84, 647–657 (2003).

14. Feist, A. M. et al. Model-driven evaluation of the production potential for growth-coupled products of *Escherichia coli*. Metab. Eng. 12, 173–186 (2010).

15. Lewis, N. E., Nagarajan, H. & Palsson, B. Ø. Constraining the metabolic genotype-phenotype relationship using a phylogeny of *in silico* methods. Nat. Rev. Microbiol. 10, 291–305 (2012).

16. Yim, H. et al. Metabolic engineering of *Escherichia coli* for direct production of 1,4-butanediol. Nat. Chem. Biol. 7, 445–452 (2011).

17. O’Brien, E. J., Lerman, J. A., Chang, R. L., Hyduke, D. R. & Palsson, B. Ø. Genome-scale models of metabolism and gene expression extend and refine growth phenotype prediction. Mol. Syst. Biol. 9, 693 (2013).

18. O’Brien, E. J. & Palsson, B. O. Computing the functional proteome: recent progress and future prospects for genome-scale models. Curr. Opin. Biotechnol. 34, 125–134 (2015).

19. Lerman, J. a. et al. In silico method for modelling metabolism and gene product expression at genome scale. Nat. Commun. 3, 929 (2012).

20. Molenaar, D., van Berlo, R., de Ridder, D. & Teusink, B. Shifts in growth strategies reflect tradeoffs in cellular economics. Mol. Syst. Biol. 5, 323 (2009).

21. Winkler, J. D., Halweg-Edwards, A. L. & Gill, R. T. The LASER database: Formalizing design rules for metabolic engineering. Metabolic Engineering Communications 2, 30–38 (2015).

22. Orth, J. D., Thiele, I. & Palsson, B. Ø. What is flux balance analysis? Nat. Biotechnol. 28, 245–248 (2010).

23. Nakahigashi, K. et al. Systematic phenome analysis of *Escherichia coli* multiple-knockout mutants reveals hidden reactions in central carbon metabolism. Mol. Syst. Biol. 5, 306 (2009).

24. Guzmán, G. I. et al. Model-driven discovery of underground metabolic functions in *Escherichia coli*. Proceedings of the National Academy of Sciences 112, 929–934 (2015).

25. Trinh, C. T., Li, J., Blanch, H. W. & Clark, D. S. Redesigning Escherichia coli metabolism for anaerobic production of isobutanol. Appl. Environ. Microbiol. 77, 4894–4904 (2011).

26. Stols, L. & Donnelly, M. I. Production of succinic acid through overexpression of NAD+-dependent malic enzyme in an *Escherichia coli* mutant. Appl. Environ. Microbiol. 63, 2695–2701 (1997).

27. Zhou, S., Shanmugam, K. T. & Ingram, L. O. Functional replacement of the *Escherichia coli* d-(−)-lactate dehydrogenase gene (ldha) with the l-(+)-lactate dehydrogenase gene (ldhl) from pediococcus acidilactici. Appl. Environ. Microbiol. 69, 2237 (2003).

28. Zhang, X., Jantama, K., Moore, J. C., Shanmugam, K. T. & Ingram, L. O. Production of l-alanine by metabolically engineered escherichia coli. Appl. Microbiol. Biotechnol. 77, 355–366 (2007).

29. Maklashina, E., Berthold, D. a. & Cecchini, G. Anaerobic expression of *Escherichia coli* succinate dehydrogenase: functional replacement of fumarate reductase in the respiratory chain during anaerobic growth. J. Bacteriol. 180, 5989–5996 (1998).

30. Ibarra, R. U., Edwards, J. S. & Palsson, B. O. *Escherichia coli* K-12 undergoes adaptive evolution to achieve in silico predicted optimal growth. Nature 420, 20–23 (2002).

31. Edwards, J. S., Ibarra, R. U. & Palsson, B. O. In silico predictions of *Escherichia coli* metabolic capabilities are consistent with experimental data. Nat. Biotechnol. 19, 125–130 (2001).

32. Ali Ebrahim, Elizabeth Brunk, Justin Tan, Edward J. O’Brien, Donghyuk Kim, Richard Szubin, Joshua A. Lerman, Anna Lechner, Anand Sastry, Aarash Bordbar, Adam M. Feist, Bernhard O. Palsson. Multi-omic data integration enables discovery of hidden biological regularities In revision.

33. Lewis, N. E. et al. Omic data from evolved *E. coli* are consistent with computed optimal growth from genome-scale models. Mol. Syst. Biol. 6, 390 (2010).

34. Atsumi, S. et al. Engineering the isobutanol biosynthetic pathway in escherichia coli by comparison of three aldehyde reductase/alcohol dehydrogenase genes. Appl. Microbiol. Biotechnol. 85, 651–657 (2010).

35. Atsumi, S., Hanai, T. & Liao, J. C. Non-fermentative pathways for synthesis of branched-chain higher alcohols as biofuels. Nature 451, 86–89 (2008).

36. Dellomonaco, C., Clomburg, J. M., Miller, E. N. & Gonzalez, R. Engineered reversal of the *β*-oxidation cycle for the synthesis of fuels and chemicals. Nature 476, 355–359 (2011).

37. McMurry, J. et al. Navigating the phenotype frontier: The monarch initiative (2016).

38. Check Hayden, E. Synthetic biologists seek standards for nascent field. Nature News 520, 141 (2015).

39. King, Z. A., Lloyd, C. J., Feist, A. M. & Palsson, B. O. Next-generation genome-scale models for metabolic engineering. Curr. Opin. Biotechnol. 35, 23–29 (2015).

40. Zhao, Q., Stettner, A. I., Reznik, E., Paschalidis, I. C. & Segrè, D. Mapping the landscape of metabolic goals of a cell. Genome Biol. 17, 109 (2016).

41. Schuetz, R., Kuepfer, L. & Sauer, U. Systematic evaluation of objective functions for predicting intracellular fluxes in *Escherichia coli*. Mol. Syst. Biol. 3, 119 (2007).

42. Lee, S. Y. Bacterial polyhydroxyalkanoates. Biotechnol. Bioeng. 49, 1–14 (1996).

43. Kartal, O., Mahlow, S., Skupin, A. & Ebenhöh, O. Carbohydrate-active enzymes exemplify entropic principles in metabolism. Mol. Syst. Biol. 7, 542 (2011).

44. King, Z. A. et al. BiGG models: A platform for integrating, standardizing and sharing genome-scale models. Nucleic Acids Res. 44, D515–22 (2016).

45. Orth, J. D. et al. A comprehensive genome-scale reconstruction of *Escherichia coli* metabolism—2011. Mol. Syst. Biol. 7, 535 (2011).

46. King, Z. A. et al. Escher: A web application for building, sharing, and embedding data-rich visualizations of biological pathways. PLoS Comput. Biol. 11, e1004321 (2015).

47. Ebrahim, A., Lerman, J. A., Palsson, B. O. & Hyduke, D. R. COBRApy: COnstraints-Based reconstruction and analysis for python. BMC Syst. Biol. 7, 74 (2013).

48. Ingram, L. O., Conway, T., Clark, D. P., Sewell, G. W. & Preston, J. F. Genetic engineering of ethanol production in *Escherichia coli*. Appl. Environ. Microbiol. 53, 2420–2425 (1987).

49. Bar-Even, A., Noor, E. & Savir, Y. The moderately efficient enzyme: Evolutionary and physicochemical trends shaping enzyme parameters. Biochemistry 4402–4410 (2011).

50. Palsson, B. Ø. Systems Biology: Properties of Reconstructed Networks (Cambridge University Press, Cambridge, UK, 2006).

51. Reed, J. L., Vo, T. D., Schilling, C. H. & Palsson, B. Ø. An expanded genome-scale model of *Escherichia coli* K-12 (iJR904 GSM/GPR). Genome Biol. 4, R54 (2003).

52. Feist, A. M. et al. A genome-scale metabolic reconstruction for *Escherichia coli* K-12 MG1655 that accounts for 1260 ORFs and thermodynamic information. Mol. Syst. Biol. 3, 121 (2007).

53. Blankschien, M. D., Clomburg, J. M. & Gonzalez, R. Metabolic engineering of *Escherichia coli* for the production of succinate from glycerol. Metab. Eng. 12, 409–419 (2010).

54. Donnelly, M. I., Millard, C. S., Clark, D. P., Chen, M. J. & Rathke, J. W. A novel fermentation pathway in an *Escherichia coli* mutant producing succinic acid, acetic acid, and ethanol. Appl. Biochem. Biotechnol. 70, 187–198 (1998).

55. Lee, S. J. et al. Metabolic engineering of *Escherichia coli* for enhanced production of succinic acid, based on genome comparison and in silico gene knockout simulation. Appl. Environ. Microbiol. 71, 7880 (2005).

56. Ma, J. et al. Enhancement of succinate production by metabolically engineered *Escherichia coli* with co-expression of nicotinic acid phosphoribosyltransferase and pyruvate carboxylase. Appl. Microbiol. Biotechnol. (2013).

57. Sánchez, A. M., Bennett, G. N. & San, K.-Y. Novel pathway engineering design of the anaerobic central metabolic pathway in *Escherichia coli* to increase succinate yield and productivity. Metab. Eng. 7, 229–239 (2005).

58. Sánchez, A. M., Bennett, G. N. & San, K.-Y. Efficient succinic acid production from glucose through overexpression of pyruvate carboxylase in an *Escherichia coli* alcohol dehydrogenase and lactate dehydrogenase mutant. Biotechnol. Prog. 21, 358–365 (2005).

59. Singh, A., Cher Soh, K., Hatzimanikatis, V. & Gill, R. T. Manipulating redox and ATP balancing for improved production of succinate in *E. coli*. Metab. Eng. 13, 76–81 (2011).

60. Stols, L., Kulkarni, G., Harris, B. G. & Donnelly, M. I. Expression of ascaris suum malic enzyme in a mutant escherichia coli allows production of succinic acid from glucose. Appl. Biochem. Biotechnol. 63–65, 153–158 (1997).

61. Vemuri, G. N., Eiteman, M. A. & Altman, E. Effects of growth mode and pyruvate carboxylase on succinic acid production by metabolically engineered strains of *Escherichia coli*. Appl. Environ. Microbiol. 68, 1715–1727 (2002).

62. Zhang, X., Shanmugam, K. T. & Ingram, L. O. Fermentation of glycerol to succinate by metabolically engineered strains of *Escherichia coli*. Appl. Environ. Microbiol. 76, 2397–2401 (2010).

